# *Arabidopsis LRR-MAL Receptor-like Kinases* regulate intraspecific and interspecific pollen-stigma interactions

**DOI:** 10.1101/2023.10.16.562574

**Authors:** Hyun Kyung Lee, Laura E. Canales Sanchez, Stephen J. Bordeleau, Daphne R. Goring

**Affiliations:** Department of Cell & Systems Biology, University of Toronto, Toronto, Canada M5S 3B2; Centre for the Analysis of Genome Evolution & Function, University of Toronto, Toronto, Canada M5S 3B2

## Abstract

Flowering plants contain tightly controlled pollen-pistil interactions required for promoting intraspecies fertilization and preventing interspecies hybridizations. In *Arabidopsis*, several receptor kinases (RKs) are known to regulate the later stages of intraspecies pollen tube growth and ovular reception in the pistil, but less is known about RK regulation of the earlier stages. The *Arabidopsis RKF1* cluster of *Leucine-Rich Repeat Malectin* (*LRR-MAL) RKs* was previously found to function in the stigma to promote intraspecies pollen hydration. Here, we tested additional combinations of up to seven *Arabidopsis LRR-MAL RK* knockout mutants for the *RKF1* cluster, *LIK1*, *RIR1* and *NILR2*. These *LRR-MAL RKs* were discovered to function in the female stigma to support intraspecies *Arabidopsis* pollen tube growth and to establish a pre-zygotic interspecies barrier against *Capsella rubella* pollen. Thus this study uncovered new biological functions for these poorly understood group of *RKs* in regulating the early stages of *Arabidopsis* sexual reproduction.

## Introduction

Sexual reproduction in flowering plants requires coordinated communication between the male pollen and the female pistil for successful fertilization (Johnson et al., 2019; Hafidh and Honys, 2021; Kim et al., 2021). This process is initiated when pollen grains first come in contact with the stigma located at the top of the pistil. Pollen can originate from an anther within the same bisexual flower or be carried by the wind or insect pollinators from other flowering plant species. In *Arabidopsis thaliana* (*Arabidopsis*), the stigma is responsible for the initial selection of intraspecific pollen and establishing a post-pollination barrier against interspecific pollen (Broz and Bedinger, 2021; Bordeleau et al., 2022; Tsuchimatsu and Fujii, 2022; Wang and Filatov, 2023). Pollen arrives in a desiccated state, and for *Arabidopsis* pollen, water is released from the stigma to allow for pollen hydration and germination. This is followed by the emergence of a pollen tube which grows directionally down the pistil, carrying two sperm cells, to an unfertilized ovule. Upon entering an ovule, the pollen tube bursts to release the sperm cells, and a double fertilization event follows with the sperm cell fusing to the female gametophyte egg cell and central cell to initiate seed development (Johnson et al., 2019; Hafidh and Honys, 2021; Kim et al., 2021). Cell-cell communication between compatible pollen and the pistil is critical for ensuring a successful outcome, but it also establishes pre-zygotic barriers to impede incompatible pollen from reaching ovules for fertilization (Broz and Bedinger, 2021; Tsuchimatsu and Fujii, 2022; Wang and Filatov, 2023).

Receptor kinases (RKs) with their respective peptide ligands play critical roles throughout this process, from the initial pollen contact with the stigma to the final stages of pollen tube reception and sperm cell release in the ovule (Johnson et al., 2019; Hafidh and Honys, 2021; Kim et al., 2021; Bordeleau et al., 2022). The large family of predicted Receptor-like Kinases (RLKs) genes in *Arabidopsis* are divided in several subgroups based on different motifs found in their predicted extracellular domains (Shiu and Bleecker, 2001; Lehti-Shiu and Shiu, 2012; Bender and Zipfel, 2023). The *Arabidopsis* RKs implicated in pollen-pistil interactions primarily include Malectin-Like (MAL-L) RKs (also known as CrRLK1Ls)(Oelmuller et al., 2023) or the Leucine-Rich Repeat (LRR) RKs that play critical roles in the later stages of pollen tube guidance and reception at an unfertilized ovule (Bordeleau et al., 2022; Yu et al., 2022). The RKs perceive a number of different secreted signals belonging to the cysteine-rich peptide (CRP) family, including LUREs and RALFs (Takeuchi, 2021; Bender and Zipfel, 2023). Two *Arabidopsis MAL-L RKs*, *FERONIA* and *ANJEA*, were also found to function in the stigma as negative regulators of pollen hydration (Liu et al., 2021), and two *Arabidopsis LRR RKs*, *PSKR1* and *PSKR2*, are required in the pistil to support pollen tube growth (Stuhrwohldt et al., 2015). Recent studies have found that *Arabidopsis* pistil factors regulating reproduction, such as *LUREs*, *TICKETs* and *FERONIA,* can not only favour intraspecific pollen tubes, but provide a barrier to closely-related interspecific pollen (Meng et al., 2019; Zhong et al., 2019; Huang et al., 2023). We previously identified predicted *Arabidopsis LRR-Malectin* (*MAL*) *RKs* in the LRR VIII-2 RK subgroup (Oelmuller et al., 2023) participating in the earlier stages of pollen-pistil interactions (Lee and Goring, 2021; Bordeleau et al., 2022). *LRR-MAL RKs* are broadly expressed in *Arabidopsis* (Lee and Goring, 2021) and diverse roles have been proposed including immune responses and perceiving cell wall-derived oligosaccharides (Le et al., 2014; Xu et al., 2014; Mendy et al., 2017; Li et al., 2020; Abel et al., 2021; Tseng et al., 2022; Martin-Dacal et al., 2023; Oelmuller et al., 2023). Our findings showed that *RECEPTOR-LIKE KINASE IN FLOWERS1* (*RKF1*) (Takahashi et al., 1998) and its tandemly-linked paralogs functioned in the stigma as positive regulators of pollen hydration (Lee and Goring, 2021). The *RKF1* cluster also functioned synergistically in the pistil with the *LRR-RKs*, *SERK1* and *BAK1*, to support pollen tube growth (Lee and Goring, 2021). In this study, we focused on the *Arabidopsis LRR-MAL RK* subgroup and identified additional *LRR-MAL RKs* that function in the pistil to support wildtype pollen. Furthermore, we uncovered a role for these *LRR-MAL RKs* in establishing an interspecies reproductive barrier.

## Results

### Reduced hydration of wildtype Col-0 pollen grains on the *LRR-MAL RK* mutant stigmas

The *Arabidopsis RKF1* gene is tandemly-linked to the *RKFL1-3* paralogs, and previously, we had used CRISPR/Cas9 to delete this gene cluster creating the *rkfΔ* mutants (Lee and Goring, 2021) (Fig. 1A). To address whether other *Arabidopsis LRR-MAL RKs* (LRR-VIII-2 RK subgroup) function with the *RKF1* cluster to regulate early pollen responses in the pistil, *LRR-MAL RK* genes expression profiles in the stigma were examined (Supplemental Table S1). *LIK1* and *RIR1* consistently displayed the highest expression across different datasets and were selected for further analysis. *NILR2* was also included as it is tandemly-linked to *RIR1* and shows good expression in the stigma as well (Supplemental Table S1). To generate mutants for these genes, CRISPR/Cas9 was used to create the *LIK1* and *NIRL2-RIR1* gene deletions in the wildtype Col-0 or the *rkfΔ-1* backgrounds (Fig. 1A, Supplemental Fig. S1, Supplemental Table S2). The null mutants were then combined with each other to create higher order mutants generating the *rkfΔ-1 nilr2rir1Δ-1 lik1-5* mutant containing seven *LRR-MAL RK* genes knocked out (Fig. 1A, Supplemental Table S2). Overall, the *rkfΔ-1 nilr2rir1Δ-1 lik1-5* septuple mutant displayed wildtype flowers and plant morphology and did not show any visible developmental or morphological defects in the pistil that could potentially impact pollen-pistil interactions (Supplemental Fig. S2). Prior evidence from our group showed that wildtype pollen hydrated more slowly on stigmas from the *rkfΔ* mutants (Lee and Goring, 2021) and so to test if the additional *LRR-MAL RK* genes were involved in this process, pollen hydration assays were conducted. For this assay, wildtype Col-0 pollen grains were placed on stigmas from wildtype Col-0 and the *LRR-MAL RK* mutant flowers, and the pollen diameters were measured as a proxy for pollen hydration(Lee et al., 2020). Decreased hydration was observed at 10 min for wildtype pollen grains on the *rkfΔ-1* stigma, but stigmas from the *rkfΔ-1 nilr2rir1Δ-1 lik1-5* septuple mutant were found to support a similar level of reduced wildtype Col-0 pollen hydration (Fig. 1B). This indicated that the *RKF1* cluster is primarily responsible in the stigma for promoting pollen hydration, and that there are no additive effects observed by knocking out three additional *LRR-MAL RK* genes.

**Figure 1.**
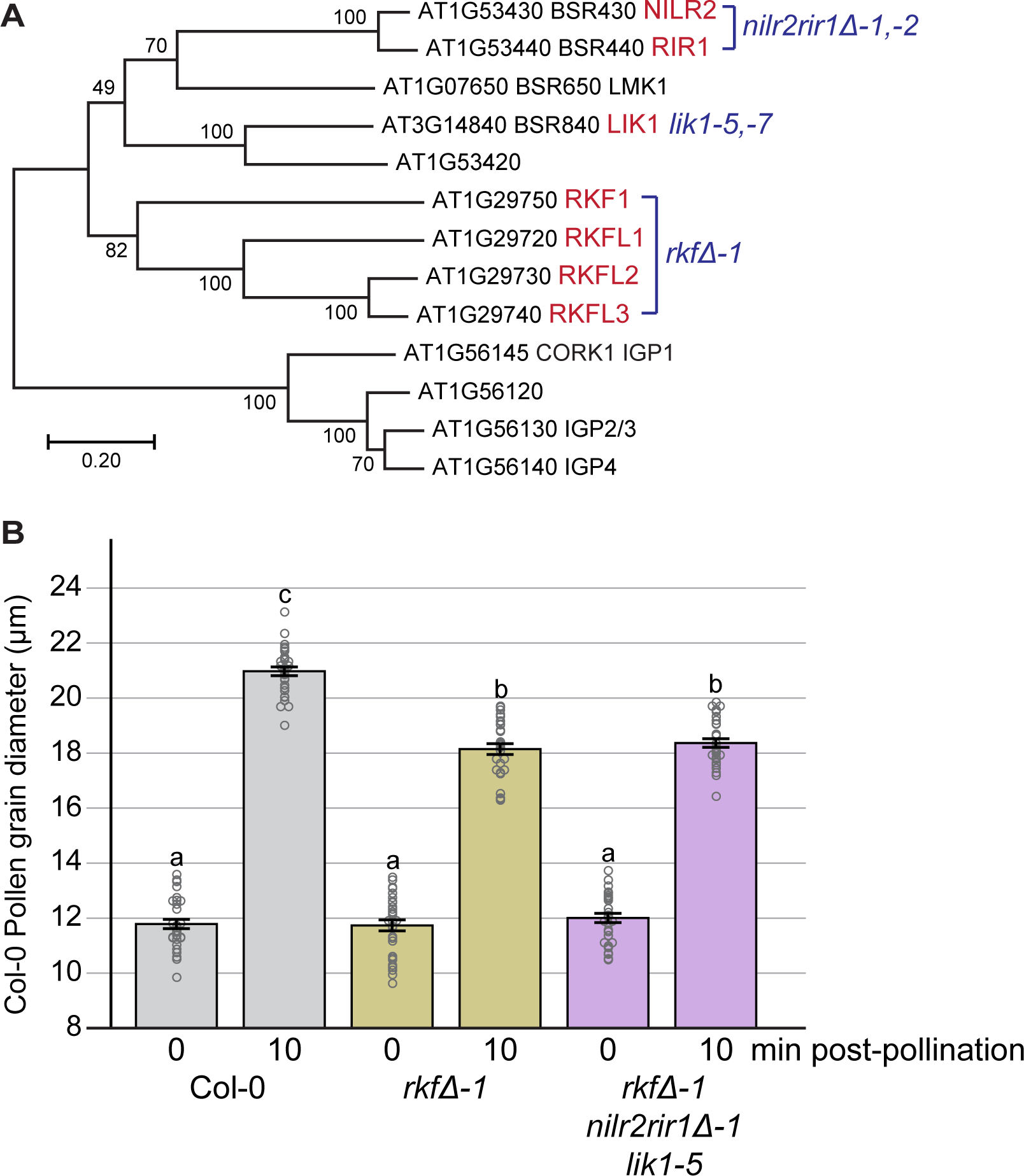
*Arabidopsis LRR-MAL RK* mutants and pollen hydration assays. **A)** Phylogenetic analysis of the predicted *Arabidopsis* LRR-MAL RK proteins (LRR-VIII-2 RK subgroup). Names in red and blue correspond to the genes and mutants, respectively, in this study. Abbreviations: BSR (BRASSINOSTEROID-SIGNALING KINASE3 (BSK3)-INTERACTING RLKs) 430, 440, 650, 840(Xu et al., 2014); NILR2 (NEMATODE-INDUCED LRR-RLK2)(Mendy et al., 2017); RIR1 (REMORIN-INTERACTING RECEPTOR1)(Abel et al., 2021); LMK1 (LEUCINE-RICH REPEAT MALECTIN KINASE1)(Li et al., 2020); LIK1 (LysM RLK1-INTERACTING KINASE1)(Le et al., 2014); RKF1 (RECEPTOR-LIKE KINASE IN FLOWERS1)(Takahashi et al., 1998); RKFL (RKF-LIKE) 1-3(Lee and Goring, 2021); CORK1 (CELLOOLIGOMER-RECEPTOR KINASE1)(Tseng et al., 2022); IGP (IMPAIRED IN GLUCAN PERCEPTION) 1, 2/3, 4(Martin-Dacal et al., 2023). **B)** Wildtype Col-0 pollen grains hydrate more slowly on stigmas from *LRR-MAL RK* mutants compared to wildtype Col-0 stigmas. All stigmas were pollinated with wildtype pollen and hydration was measured by taking pollen grain diameters at 0- and 10-minutes post-pollination. n = 30 pollen grains per line. Data is shown as a bar graph of means ± SE with all data points displayed. Letters represent statistically significant groupings of P<0.05 based on a one-way ANOVA with a Tukey-HSD post-hoc test.

### Patterns of wildtype Col-0 pollen tube growth in the *LRR-MAL RK* mutant pistils

While the wildtype Col-0 pollen grains were less hydrated at 10 min post-pollination on the *LRR-MAL RK* mutant stigmas, this phenotype was not sufficient to block pollen germination and the emergence of pollen tubes (Rozier et al., 2020). To next assess if pollen tube growth was impacted through the stigma, *LRR-MAL RK* mutant pistils were pollinated with wildtype Col-0 pollen grains and harvested at 2-hours post-pollination. The pollen tubes were visualized by aniline blue staining which stains for callose, a ß-1,3 glucan that is deposited both in the pollen tube cell wall and as plugs which separate the cytoplasm in the growing pollen tube tip from the remainder of the pollen tube (Franklin-Tong, 1999; Chebli et al., 2012). At 2-hours post-pollination, the wildtype Col-0 pollen tubes have largely grown through the stigma and style, and entered into the transmitting tract for all the different lines (Fig. 2, Supplemental Fig. S3). Interestingly though, the callose plugs appeared to take on a different shape in pollen tubes growing through some of the *LRR-MAL RK* mutant stigmas (Fig. 2, Supplemental Fig. S3). In the wildtype Col-0 pistils, the callose plugs in the pollen tubes were the typical narrow and elongated shape (Fig 2C) while they appeared to be shorter and wider in the wildtype pollen tubes growing through the *nilr2rir1Δ-1 lik1-5* and *rkfΔ-1 nilr2rir1Δ-1 lik1-5* mutant stigmas (Fig 2F, I) but not for pistils from the other *LRR-MAL RK* mutant combinations (Supplemental Fig. S3). By measuring the length, width, and area of the callose plugs, the initial visual observations were supported as the pollen tube callose plugs in both the *nilr2rir1Δ-1 lik1-5* and *rkfΔ-1 nilr2rir1Δ-1 lik1-5* mutant stigmas displayed significantly reduced lengths, increased widths, and smaller areas (Fig 2N-P). As well the pollen tube callose plugs in the *rkfΔ-1 nilr2rir1Δ-1 lik1-5* septuple mutant stigmas displayed a further decrease in the length and increase in width compared to those in the *nilr2rir1Δ-1 lik1-5* triple mutant stigmas signifying an additive effect when all seven *LRR-MAL RK* genes were knocked out in the stigma. This altered callose plug phenotype was largely rescued by the stigma-specific expression of *RKF1* (*SLR1:RKF1* transgene) in the *rkfΔ-1 nilr2rir1Δ-1 lik1-5* mutant (Fig 2N-P). Overall, the change observed in the wildtype pollen tube callose plugs when growing through the *LRR-MAL RK* mutant stigmas is a novel phenotype that has not been reported before. The increased callose plug width may be indicative of wider wildtype pollen tubes resulting from changes in the growth environment (Reimann et al., 2020; Kapoor and Geitmann, 2023) of the *LRR-MAL RK* mutant stigmas.

**Figure 2.**
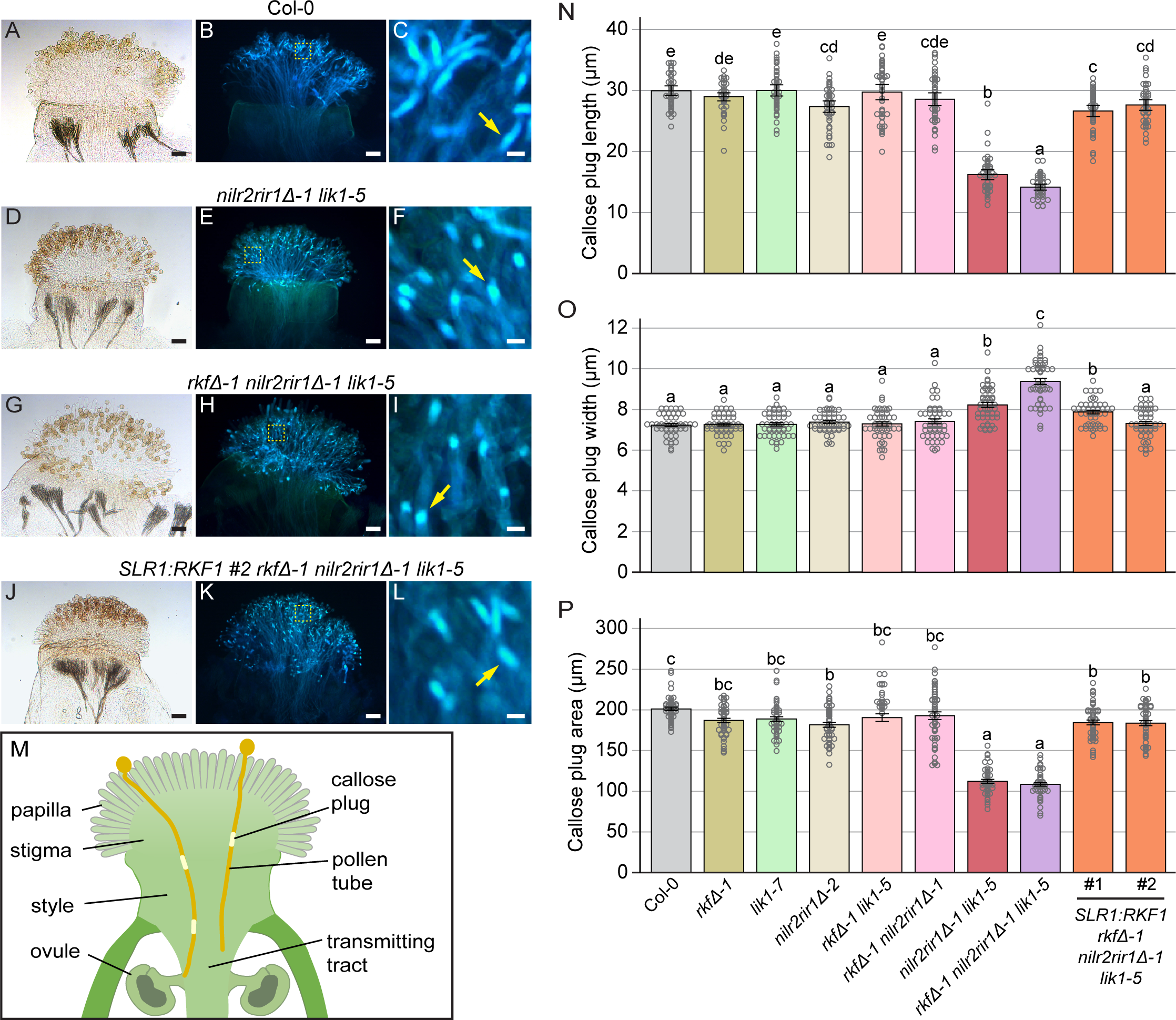
Altered callose plug sizes in *Arabidopsis* wildtype Col-0 pollen tubes growing through *LRR-MAL RK* mutant stigmas at 2 hrs post-pollination. **A-L)** Representative images of aniline blue stained pistils from wildtype Col-0, *LRR-MAL RK* mutants, and a *SLR1:RKF1* rescue line. All stigmas were pollinated with wildtype pollen and harvested for aniline-blue staining at 2-hours post-pollination. For each line, brightfield images are shown on the left and aniline blue stained images on the right. The *SLR1:RKF1* transgene drives the stigma-specific expression of *RKF1* to rescue the callose plug phenotype in the wildtype pollen tubes (**k-l**). Dashed yellow boxes in **b,e,h,k** outline the regions shown in **c,f,i,l**. Yellow arrows in **c,f,i,l** point to an example of a callose plug. Scale bar for **a-b, d-e, g-h, j-k** = 100 µm; Scale bar for **c,f,i,l** = 500 µm. See Supplemental Fig. 4 for representative images for all the different *LRR-MAL RK* mutant pistils. **M)** Model illustrating callose plugs in pollen tubes growing through the reproductive tract to an ovule. **N-P)** Bar graphs displaying the length, width and area of callose plugs in the wildtype pollen tubes growing through the stigmas/styles from wildtype Col-0 and the *LRR-MAL RK* mutant pistils as indicated at the bottom of the graphs. Data are shown as means ± SE with all data points displayed. n = 50 callose plugs per line. Letters represent statistically significant groupings of P<0.05 based on a one-way ANOVA with a Tukey-HSD post-hoc test.

Previously, we used wildtype Col-0 pollen carrying the *LAT52:GUS* reporter for GUS staining to quantify pollen tube fronts(Swanson et al., 2016) and found that the *rkfΔ-1* mutant pistils supported wildtype pollen tube growth at 2 and 6-hours post-pollination (Lee and Goring, 2021). However, when *rkfΔ-1* was combined with the *serk1-1 bak1-4* mutations, there was a synergistic effect in reducing wildtype pollen tube growth distances at 6-hours in the mutant pistils (Lee and Goring, 2021). Here we examined whether the wildtype Col-0 *LAT52:GUS* pollen tube growth distances were affected in pistils from the different *LRR-MAL RK* mutant combinations. Since the altered callose plug phenotype was apparent in pollen tubes growing through the *nilr2rir1Δ-1 lik1-5* and *rkfΔ-1 nilr2rir1Δ-1 lik1-5* mutant stigmas at 2 hrs post-pollination, we examined pollen tube growth fronts at this stage and did not find any significant differences compared to wildtype Col-0 pistils (Fig. 3). Similarly, the *SLR1:RKF1 rkfΔ-1 nilr2rir1Δ-1 lik1-5* pistils supported wildtype Col-0 *LAT52:GUS* pollen tube growth distances (Fig. 3). Finally, wildtype levels of seed production were observed in the *rkfΔ-1 nilr2rir1Δ-1 lik1-5* mutant and the *SLR1:RKF1 rkfΔ-1 nilr2rir1Δ-1 lik1-5* lines (Supplemental Fig. S4). Thus, the altered phenotype of callose plugs in the wildtype pollen tubes was largely confined to the stigmas from the *nilr2rir1Δ-1 lik1-5* and *rkfΔ-1 nilr2rir1Δ-1 lik1-5* mutants and did not affect downstream pollen tube growth and fertilization.

**Figure 3.**
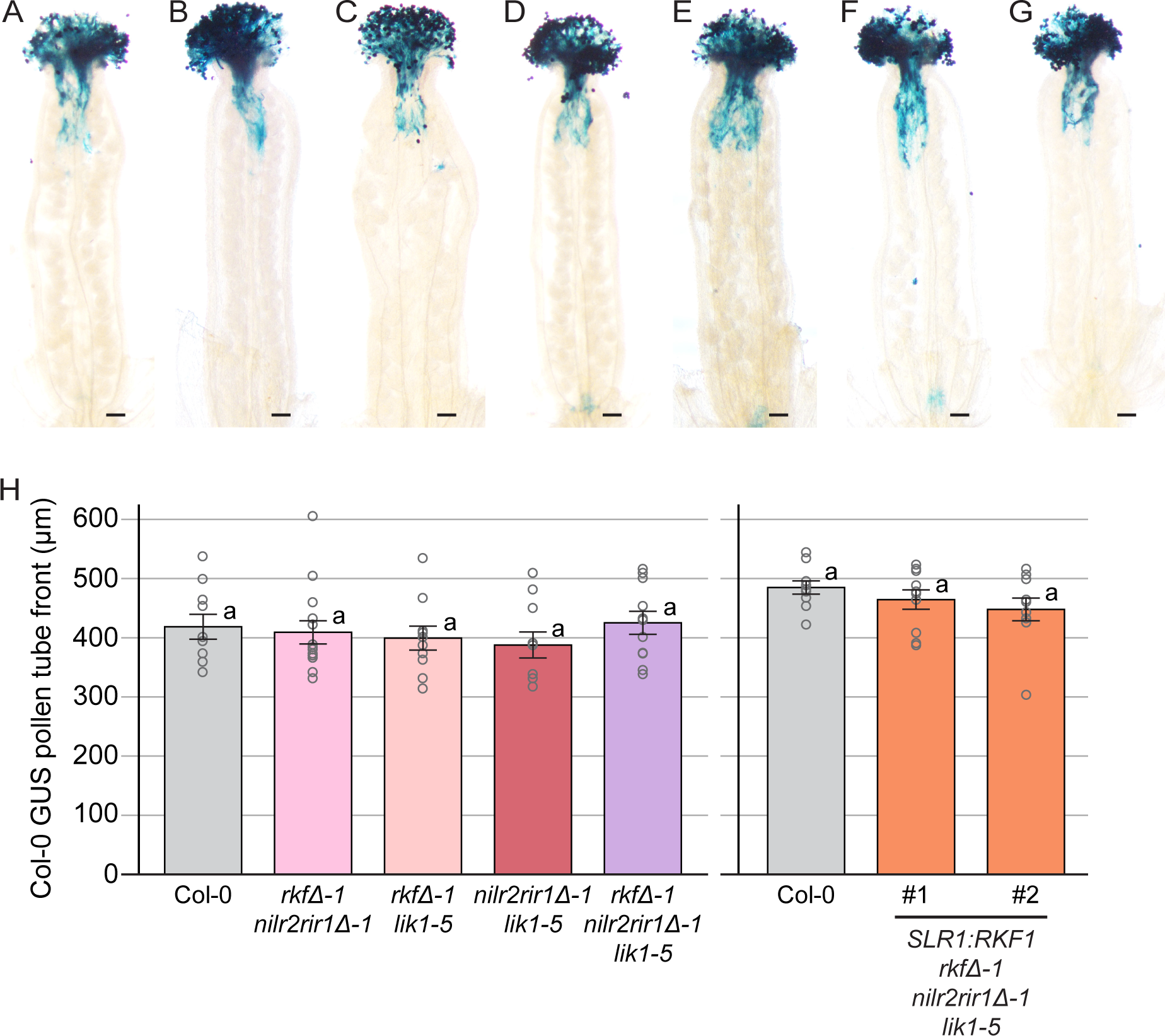
*Arabidopsis* wildtype Col-0 *LAT52:GUS* pollen tubes display normal travel distances in *LRR-MAL RK* mutant pistils at 2 hrs post-pollination. **A-G)** Representative images of GUS stained pistils from wildtype Col-0, *LRR-MAL RK* mutants, and the *SLR1:RKF1* rescue lines. All stigmas were pollinated with wildtype *LAT52:GUS* pollen and harvested for GUS staining at 2-hours post-pollination. Scale bar = 100 µm. **H)** Bar graphs displaying the lengths of the wildtype *LAT52:GUS* pollen tube fronts in pistils from wildtype Col-0, the different *LRR-MAL RK* mutants, and the *SLR1:RKF1* rescue lines. Each pollen tube front was measured from the base of the style. Data are shown as mean ± SE with all data points displayed. n = 10-14 pistils per line. No significant differences were observed based on a one-way ANOVA with a Tukey-HSD post-hoc test (P<0.05).

### *LRR-MAL RKs* establish an interspecies reproductive barrier in the stigma

Since signaling proteins involved in intraspecific pollen-pistil interactions can also have additional roles in promoting self pollen over pollen from related species (Escobar-Restrepo et al., 2007; Zhong et al., 2019), we investigated if the *Arabidopsis LRR-MAL RKs* could have a similar dual function in the stigma. Interspecific pollinations were conducted with another Brassicaceae species, the closely-related *Capsella rubella*, whose pollen had been previously shown to be incompatible with wildtype *Arabidopsis* Col-0 pistils (Fujii et al., 2019). The *Arabidopsis spri1-1* mutant, identified in a screen for factors regulating interspecies barriers (Fujii et al., 2019), was used as a positive control. *SPRI1* is a stigma-specific gene encoding a small multipass transmembrane protein that is required to reject *C. rubella* pollen tubes (Fujii et al., 2019; Kato et al., 2023). To test for changes to the interspecies barrier in the *LRR-MAL RK* mutant stigmas, pistils were harvested at 24-hours post-pollination and aniline blue stained to visualize the *C. rubella* pollen tubes. In general for all pollinations, the *C. rubella* pollen grains were found to germinate and pollen tubes could be observed at the top of the stigma (Fig. 4B,D,F,H,J,L). The influence of the interspecies barrier on *C. rubella* pollen tube growth was most apparent in how many pollen tubes successfully entered into the style. For both the *Arabidopsis spri1-1* and the *rkfΔ-1 nilr2rir1Δ-1 lik1-5* septuple mutants, significantly more *C. rubella* pollen tubes were present in the styles compared to wildtype *Arabidopsis* Col-0 pistils (Fig. 4M). In contrast, the *C. rubella* pollinated *rkfΔ-1* mutant pistils were the same as wildtype Col-0 pistils. Interestingly, not only did the stigma-specific expression of *RKF1* (*SLR1:RKF1* transgene) in the *rkfΔ-1 nilr2rir1Δ-1 lik1-5* mutant rescue the breakdown in the interspecies barrier, but in fact these lines displayed a better barrier to *C. rubella* pollen tubes with significantly fewer *C. rubella* pollen tubes present in the style (Fig. 4M). We also noticed a difference in the number of *C. rubella* pollen grains that were visible in the brightfield images of the pollinated wildtype and mutant *Arabidopsis* stigmas (Fig. 4A,C,E,G,I,K). *C. rubella* pollen grains that did not successfully germinate likely washed away during the aniline blue staining process as they may not have been well-adhered to the stigma. For both the *Arabidopsis spri1-1* and *rkfΔ-1 nilr2rir1Δ-1 lik1-5* mutant stigmas, significantly more *C. rubella* pollen grains were present compared to wildtype *Arabidopsis* Col-0 stigmas (Fig. 4N). In a similar trend to that seen for the pollen tubes, fewer *C. rubella* pollen grains were present on stigmas from the *SLR1:RKF1 rkfΔ-1 nilr2rir1Δ-1 lik1-5* rescue lines (Fig. 4N). A comparison of the ratios of the number of *C. rubella* pollen grains over pollen tubes showed no significant differences between wildtype Col-0, *spri1-1*, *rkfΔ-1* and *rkfΔ-1 nilr2rir1Δ-1 lik1-5* pistils. This result points to a barrier that appears to be impacting the earlier stage of interspecies pollen germination which in turn is reflected in the number of *C. rubella* pollen tubes that successfully enter the style (Fig. 4O). Interestingly, this analysis also uncovered a significantly larger ratio for the *SLR1:RKF1 rkfΔ-1 nilr2rir1Δ-1 lik1-5* rescue lines which indicates that the *SLR1* promoter driven expression of *RKF1* in the stigma is having a stronger impact in blocking *C. rubella* pollen tubes from entering into the style. Overall, these results show that the *LRR-MAL RKs* have a function in the stigma to establish a post-pollination barrier to restricts interspecies pollen tube growth.

**Figure 4.**
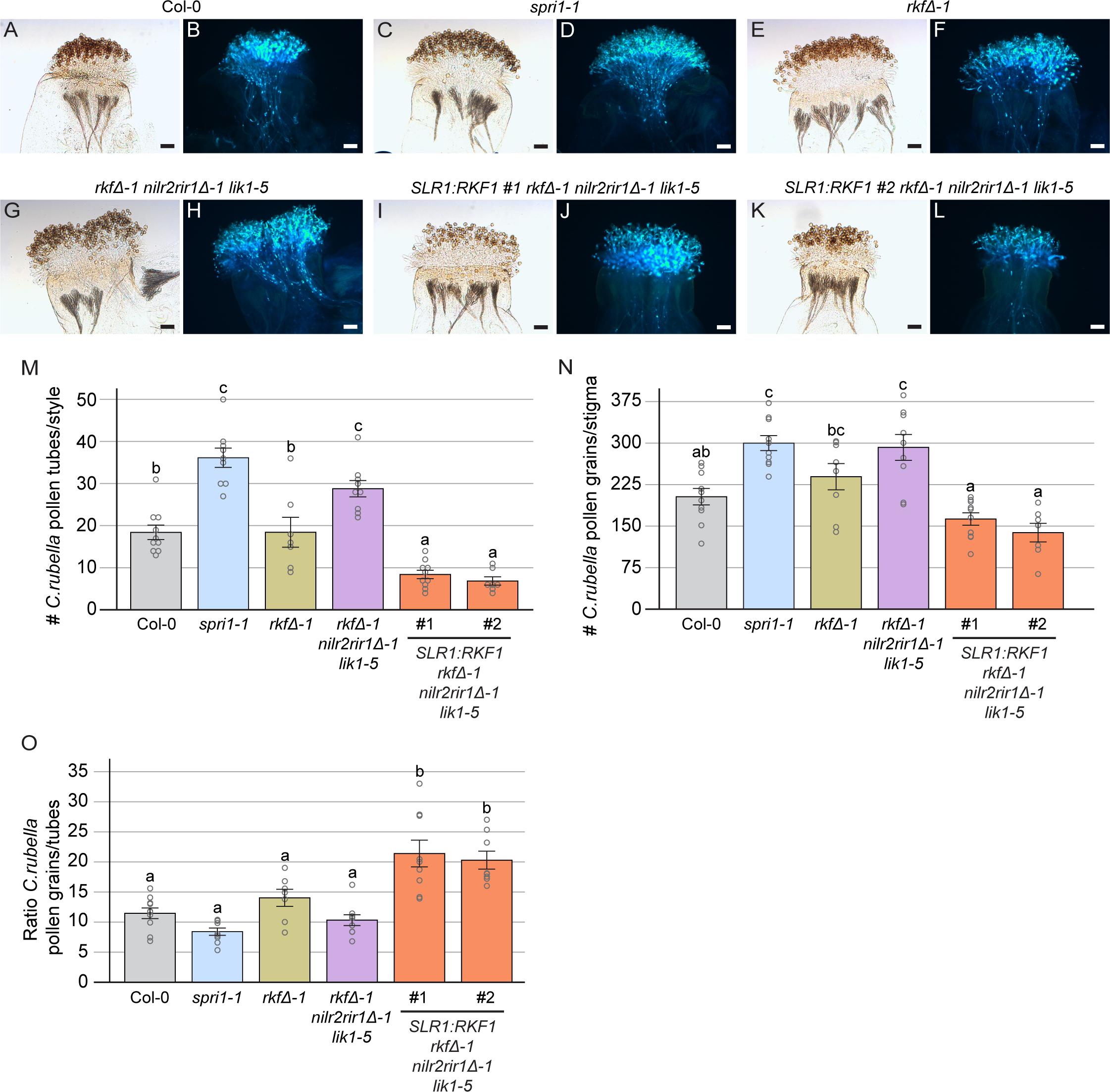
A breakdown in interspecies incompatibility is seen when *Arabidopsis LRR-MAL RK* mutant stigmas are pollinated with *Capsella rubella* pollen. **A-L)** Representative images of aniline blue stained pistils from *Arabidopsis* wildtype Col-0, *spri1-1*, *rkfΔ-1*, *rkfΔ-1 nilr2rir1Δ-1 lik1-5* and *SLR1:RKF1 #2 rkfΔ-1 nilr2rir1Δ-1 lik1-5* flowers. All stigmas were pollinated with *C. rubella* pollen and stained at 24-hours post-pollination. For each line, brightfield images are shown on the left and aniline blue stained images on the right. The *spri1-1* mutant was used as a positive control for interspecies incompatibility breakdown(Fujii et al., 2019). Scale bar = 100 µm. **M-O)** Bar graphs displaying the number of *C. rubella* pollen tubes/style (**m**) and *C. rubella* pollen grains/stigma (**n**) on *Arabidopsis* wildtype Col-0 and mutant pistils, and the ratios of these two data sets (# pollen grains/pollen tubes) (**o**). Data are shown as means ± SE with all data points displayed. n = 7-10 pistils per line. Letters represent statistically significant groupings of P<0.05 based on a one-way ANOVA with a Tukey-HSD post-hoc test.

## Discussion

Several different *Arabidopsis* RKs are known to play critical roles in regulating pollen-pistil interactions, yet little is known about RKs that function in the stigma/style region of the pistil to promote compatible pollen-pistil interactions (Kim et al., 2021; Bordeleau et al., 2022).

Previously, we had uncovered a role for one set of *Arabidopsis LRR-MAL RKs,* the *RKF1* cluster, in the stigma as positive regulators of pollen hydration (Lee and Goring, 2021). Furthermore, the *RKF1* cluster in combination with two *LRR RKs*, *SERK1* and *BAK1*, was found to promote wildtype pollen tube growth in the reproductive tract. While pistils from the *RKF1* cluster mutant (*rkfΔ*) displayed normal *Arabidopsis* pollen tube growth, pistils from the *serk1 bak1* double mutant supported reduced wildtype pollen tube growth. However, wildtype pollen tube growth was further reduced in pistils from the combined *rkfΔ serk1 bak1* mutant indicating a synergistic effect between these two groups of *Arabidopsis RKs*. In this study, we exclusively focused on the *LRR-MAL RKs* and uncovered specific functions in the stigma/style for seven *LRR-MAL RKs* in supporting intraspecific pollen over interspecific pollen.

In addition to the previously defined role of the *RKF1* cluster in the stigma to support pollen hydration (Fig. 1), the *RKF1* cluster along with *NILR2*, *RIR1* and *LIK1* are required in the stigma to support wildtype pollen tubes. In the *rkfΔ-1 nilr2rir1Δ-1 lik1-5* septuple mutant stigmas, wildtype pollen tubes displayed callose plugs that were shorter in length and wider in diameter which is likely indicative of the pollen tubes having an increased diameter, and this phenotype was rescued by the expressing *RKF1* under the control of the stigma-specific *SLR1* promoter (Fig. 2). Pollen tubes undergo rapid polarized growth, and callose plugs are deposited to maintain turgor pressure and cytoplasm volume in the growing tip region by separating it from the older parts of the pollen tube (Kapoor and Geitmann, 2022; Kapoor and Geitmann, 2023). The *Arabidopsis* stigma along with the style are solid tissues where the pollen tubes undergo invasive growth before entering into the transmitting tract where the cells are more loosely organized (Ndinyanka Fabrice et al., 2017; Reimann et al., 2020; Kapoor and Geitmann, 2022). The wildtype pollen tubes may be changing shape as they grow through a less responsive growth environment in the *LRR-MAL RK* mutant stigmas, though there was no significant impact on pollen tube growth distances at 2-hours post-pollination (Fig. 3).

We also discovered that the *Arabidopsis LRR-MAL RKs* function in presenting a barrier in the stigma to block interspecific pollen tubes. Presenting a barrier to pollen from other species is an important trait as it allows the pistil to control mate selection and increase the rate of successful fertilization (Broz and Bedinger, 2021; Tsuchimatsu and Fujii, 2022; Wang and Filatov, 2023). For all lines tested, *C. rubella* pollen grains were able to germinate and form pollen tubes on the stigma surface, but there were significant differences in the number of *C. rubella* pollen grains on the stigma surfaces suggesting that the rate of *C. rubella* pollen grain adherence and germination was impacted (with poorly adhered pollen grains washing away during staining, Fig. 4). As well, there were significant differences in the number of *C. rubella* pollen tubes that had entered the style. Comparing the *rkfΔ-1 nilr2rir1Δ-1 lik1-5* septuple mutant to wildtype Col-0, there were significantly more *C. rubella* pollen grains present on the mutant stigma surface and *C. rubella* pollen tubes growing through the mutant styles. The breakdown of this interspecies block was comparable to the *spri1-1* mutant (Fig 4), which was previously shown to be required in the stigma to establish an interspecies barrier (Kato et al., 2023). Interestingly, the stigma-specific expression of *RKF1* by the *SLR1* promoter in the *rkfΔ-1 nilr2rir1Δ-1 lik1-5* septuple mutant not only rescued this phenotype, but these lines were in fact even more effective in rejecting *C. rubella* pollen tubes (Fig. 4).

*LRR-MAL RKs* are broadly expressed in *Arabidopsis* (Lee and Goring, 2021) and potentially have very diverse functions (Le et al., 2014; Xu et al., 2014; Mendy et al., 2017; Li et al., 2020; Abel et al., 2021; Tseng et al., 2022; Martin-Dacal et al., 2023; Oelmuller et al., 2023). It is not unusual for RKs involved in sexual reproduction to be involved in other biological processes. For example, *Arabidopsis FERONIA* was first discovered in genetic screens for female mutants affected in pollen tube reception by the female gametophyte (Huck et al., 2003; Rotman et al., 2003), and subsequently, other very diverse roles were uncovered for FERONIA including plant immunity (Zhu et al., 2021; Bender and Zipfel, 2023; Malivert and Hamant, 2023). While much less is known about *Arabidopsis* LRR-MAL RKs, some of the members have been implicated in perceiving cell wall-derived oligosaccharides and immune responses (CORK1/IGP1, IGP2/3, IGP4) (Tseng et al., 2022; Martin-Dacal et al., 2023; Oelmuller et al., 2023). For CORK1, two conserved phenylalanine residues in the MAL domain were shown to be important for perceiving cellooligomers (Tseng et al., 2022). This raises the idea that the RKF1 cluster, NILR2, RIR1 and LIK1 in the stigma and style could be perceiving cell wall-derived oligosaccharides, potentially generated by pollen tubes growing invasively through these tissues. The extracellular domains of these predicted RKs also have a LRR domain composed of 10-11 LRR repeats that is N-terminal to the MAL domain, and this LRR domain could be perceiving additional extracellular signals such as pollen-derived peptide signals (Takeuchi, 2021; Yu et al., 2022). It may be that that the combination of these two domains allow these *Arabidopsis* LRR-MAL RKs to discriminate between intraspecific and interspecific pollen tubes in the stigma. These members are predicted to encode functional kinases and kinase activity has been previously demonstrated for RKF1 and LIK1 (Takahashi et al., 1998; Le et al., 2014). The stronger interspecies incompatibility phenotype seen in the *SLR1:RKF1* rescue lines does suggest a more active role for the LRR-MAL RKs in blocking *C. rubella* pollen tubes. The loss of these LRR-MAL RKs in the stigma would then remove this barrier allowing more *C. rubella* pollen tubes to enter into the style. In terms of *Arabidopsis* pollinations, the potential increase in wildtype pollen tube diameter (as reflected by the callose plugs) growing through the *rkfΔ-1 nilr2rir1Δ-1 lik1-5* mutant stigma may indicate that the LRR-MAL RKs are needed in the stigma to support pollen tube growth. The loss of these LRR-MAL RKs in the stigma could result in a less responsive environment for intraspecies pollen tubes which causes the pollen tubes to increase in diameter to invasively grow through the tissue (Reimann et al., 2020). Overall in this study, we have identified a dual role for *LRR-MAL RKs* to preferential promote intraspecific pollen tubes over interspecific pollen tubes.

## Materials and Methods

### Plant material and growth conditions

*Arabidopsis* and *C. rubella* seeds were sterilized and stratified for 2 days and transferred to soil or first germinated on ½ MS Sucrose plates prior to transfer. The soil was supplemented with 1g/L of Plant Prod All Purpose Fertilizer 20-20-20, and plants were grown in a 22°C chamber on a 16-h light/8-h dark cycle and fertilized on a weekly basis. Seeds for *spril1-1* (SALK_047439C) and were ordered from the ABRC. All mutants were genotyped using primers listed in Supplemental Table S3.

### Phylogenetic Analysis and Expression Profiling

For the phylogenetic analysis of predicted *Arabidopsis* LRR-MAL RK proteins, amino acid sequences for the LRR-VIII-2 RK subgroup (Shiu and Bleecker, 2001; Lehti-Shiu and Shiu, 2012) were retrieved from TAIR (Araport11). In MEGA11 (Tamura et al., 2021), MUSCLE (Edgar, 2004) was set at default parameters to align the amino acid sequences, and a Maximum Likelihood Tree (Jones et al., 1992) was built using default parameters with 1000 bootstrap replications. *LRR-MAL RK* expression profiles were searched in the TRAVA transcriptome database (Klepikova et al., 2016) and two other stigma transcriptome datasets (Iwano et al., 2014; Gao et al., 2018) as well as a large scale *LRR-RK* promoter GUS activity study (Wu et al., 2016), and heat maps were generated using the BAR Heat mapper plus tool (Toufighi et al., 2005) (Supplemental Table S1).

### Vector construction, CRISPR-generated mutations and *Arabidopsis* transformation

For the *SLR1:RKF1* construct, the full length RKF1 coding region was amplified from a plasmid (Lee and Goring, 2021) using *RKF1*-attB primers and the NEB Phusion® High-Fidelity DNA polymerase. The amplified PCR product gel-extracted and gateway cloned into the pDONR207 entry clone with the BP Clonase II Enzyme Mix (ThermoFisher Scientific). Using the resulting pDONR207-RKF1 entry vector, *RKF1* was then gateway cloned into the pORE3-SLR1 destination vector with the LR Clonase II Enzyme Mix (ThermoFisher Scientific).

*Agrobacterium tumefaciens* (strain GV3101) competent cells were transformed with the pORE3-SLR1:RKF1 plasmid, and then used to transform the *Arabidopsis rkfΔ-1 lik1-5 nilr2rir1Δ-1* mutant by the floral dip method (Clough and Bent, 1998). T1 seeds were collected from the dipped *rkfΔ-1 lik1-5 nilr2rir1Δ-1* plants, sterilized, and germinated on soil were selected for by spraying with Basta. Surviving transgenic seedlings were confirmed as transformed using Basta primers and grown to flowering for further analysis. Expression of *RKF1* in the *SLR1:RKF1* rescue lines was confirmed using RT-PCR on RNA extracted from half pistils (Supplemental Fig. S1E). Approximately 30 half pistils were flash frozen and ground in liquid nitrogen, and the RNA was extracted with the Qiagen RNeasy Plant Mini Kit. To remove any gDNA contamination, a DNase treatment was performed using the Promega RQ1 RNase-Free DNase kit. Using the DNase treated RNA, cDNA was synthesized using the ThermoFisher SuperScript IV Reverse Transcriptase kit and then used in PCR reaction to amplify the RKF1 cDNA. A minus Reverse Transcriptase reaction (-RT) was also set up as a negative control for the PCR reactions. For generating deletion mutations in *NILR2-RIR1* and *LIK1*, two guide RNAs for each deletion were cloned into the pBEE401E CRISPR/Cas9 vector as previously described (Wang et al., 2015; Doucet et al., 2019; Doucet et al., 2019) and transformed into *Arabidopsis* as described above. Basta resistant transgenic T1 seedings were selected genotyped by PCR to identify deletion mutations as shown in Supplemental Fig. S1. All primers used in are listed in Supplemental Table S3.

### Pollination Assays

For all assays, *Arabidopsis* Stage 12 flower buds were emasculated in the mornings under a dissecting microscope by removing the anthers and petals using forceps. Emasculated *Arabidopsis* pistils were left on the plants and wrapped in plastic wrap with care to avoid damaging the stigmatic papillae and left overnight in the growth chamber. The next day, the plastic wrap was carefully removed to uncover a now stage 13 pistil ready for pollination. One mature anther from wildtype Col-0 or *C. rubella* was used to lightly apply a monolayer of pollen to the stigmatic surface. Pollen hydration assays were performed following the protocol outlined in (Lee et al., 2020), and images were collected using a Nikon sMz800 microscope and the NIS-elements imaging software. For pollen tube growth assays, the *Arabidopsis*-pollinated *Arabidopsis* pistils were left on the plant and harvested at 2 hours post-pollination while the *C. rubella*-pollinated *Arabidopsis* pistils were left on the plant and harvested at 24 hours post-pollination. For GUS staining of pollen tubes, pistils pollinated with wildtype Col-0 *LAT52:GUS* (Swanson et al., 2016) were fixed and stained for GUS activity as previously described (Johnson et al., 2004; Lee and Goring, 2021), and images were collected using a Nikon sMz800 microscope. Using the NIS-elements imaging software, pollen tube growth was measured by drawing a line across the bottom of the style and measuring the distance to the pollen tube front. For aniline blue staining of pollen tubes, the harvested pistils were fixed and stained with aniline blue as described in Lee et al (Lee et al., 2020), and images were taken on a Zeiss Axioskop2Plus fluorescence microscope under brightfield and UV fluorescence filter. Using the INFINITY Analyze software, the area of the callose plugs in the pollen tubes was measured by using the polygonal tool to trace a callose plug at 400X. The same software was used to measure length and width using the length tool (5 stigmas/line with 10 callose plugs/stigma). To count the number of pollen tubes/style for the *C. rubella*-pollinated *Arabidopsis* pistils, the counting tool in the NIS-elements imaging software was used. To count the number of pollen grains/stigma for the *C. rubella*-pollinated *Arabidopsis* pistils, the cell counter plugin in ImageJ was used. For seed set analysis, naturally pollinated siliques were collected and placed in an Eppendorf tube containing 70% ethanol and left for a week or until the siliques were clear (Beuder et al., 2020). Cleared siliques were mounted on a slide with water, and seeds were counted using a Nikon sMz800 microscope and the NIS-elements imaging software.

## Acknowledgements

We thank Safa Abdulsalam, Carmela Serio, Minyoung Chung and Shreya Das for technical assistance, and members of the Goring lab for critically reading this article. We are very grateful to Robert Swanson (Valparaiso U) for the Col-0 *LAT52:GUS* seeds and Stephen Wright (U Toronto) for the *C. rubella* seeds.

## Author contributions

HKL, LECS, SJB, DRG conceived and designed the experiments; HKL, LECS, SJB performed the experiments; HKL, LECS, SJB, DRG analyzed the data; HKL, LECS, DRG wrote the paper; HKL, LECS, SJB, DRG edited the paper.

## Supplemental Data

**Supplemental Figure S1. |** *nilr2rir1* and *lik1* mutants and the *SLR1:RKF1 rkfΔ-1 nilr2rir1Δ-1 lik1-5* lines.

**Supplemental Figure S2. |** Representative images of *Arabidopsis* wildtype Col-0 and *LRR-MAL RK* mutant flowering plants and flowers.

**Supplemental Figure S3. |** Representative images of aniline blue stained Col-0-pollinated pistils from *Arabidopsis* wildtype Col-0 and the *LRR-MAL RK* mutant plants.

**Supplemental Figure S4. |** Seed set on naturally pollinated pistils.

**Supplemental Table S1 |** Expression profiles for LRR-MAL RKs in stigma transcriptome datasets.

**Supplemental Table S2 |** Mutants used in this study.

**Supplemental Table S3 |** List of primers used.

