## Supplementary material for "*Arabidopsis LRR-MAL Receptor-like Kinases* regulate intraspecific and interspecific pollen-stigma interactions": Lee et al 2023 Suppl Files (from another version of this preprint: version 1, 2023-10-17)

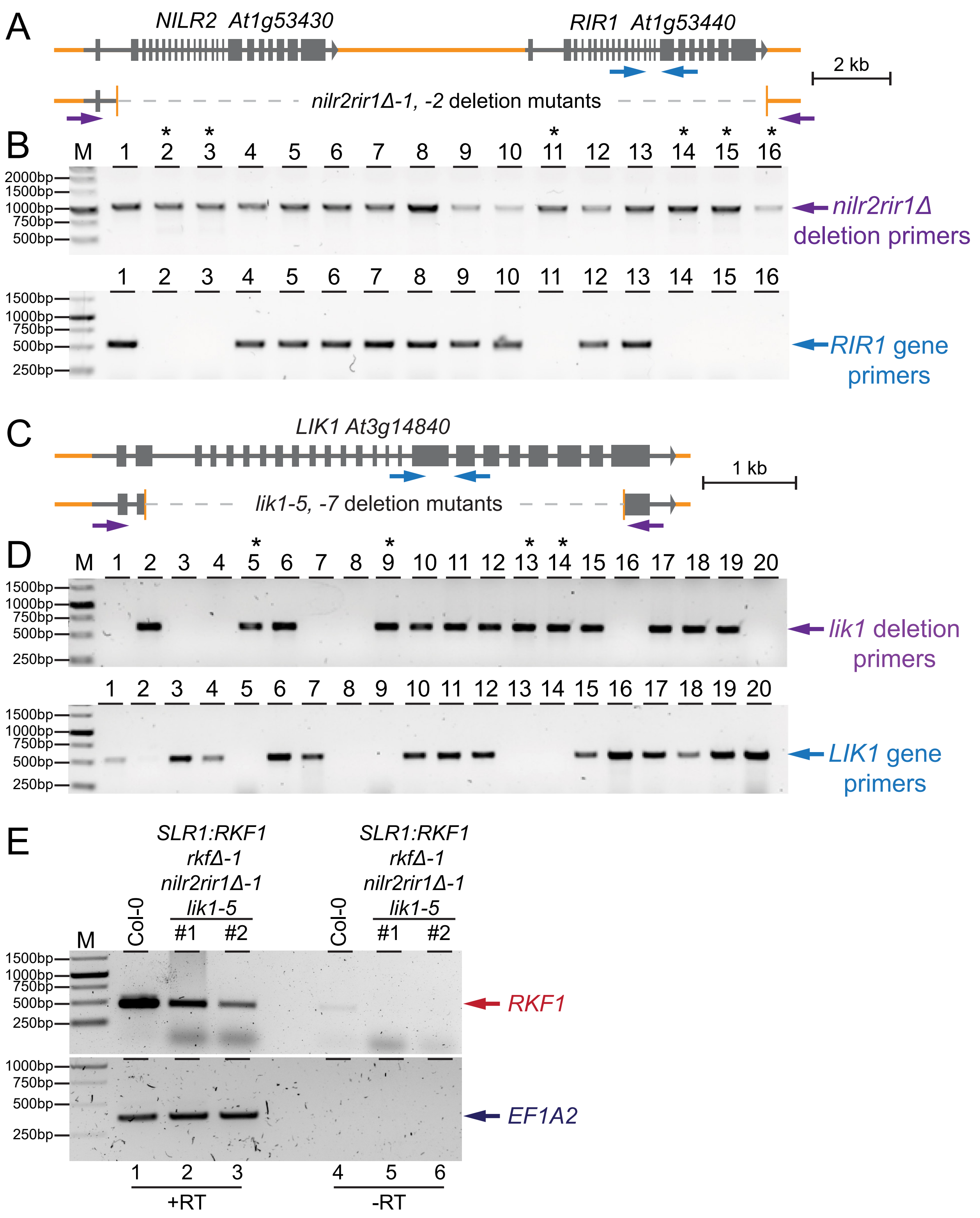


**Supplemental Figure 1. *nilr2rir1* and *lik1* mutants and the *SLR1:RKF1 rkfΔ-1 nilr2rir1Δ-1 lik1-5* lines.**
**A-B)** Generation of the CRISPR/Cas9 deletion mutations for *NILR2-RIR1*. **a**, Schematic representation of the *NILR2* and *RIR1* genes and the *nilr2rir1Δ-1, -2* deletion mutation. Locations of deletion and internal primers are indicated by the purple and blue arrows, respectively. **b**, Representative gel photos detecting the *nilr2-rir1Δ* deletion (top) and the *RIR1* gene (bottom). *mark homozygous *nilr2-rir1Δ* mutants.
**C-D)** Generation of the CRISPR/Cas9 deletion mutations for *LIK1*. **c**, Schematic representation of the *LIK1* gene and *lik1-5, -7* deletion mutation. Locations of deletion and internal primers are indicated by the purple and blue arrows, respectively. **d**, Representative gel photos detecting the *lik1* deletion (top) and the *LIK1* gene (bottom). *mark homozygous *lik1* mutants.
**E)** RT-PCR analysis of *RKF1* expression in ½ pistil RNA from Col-0 (Lane 1) and the *SLR1:RKF1 rkfΔ-1 nilr2rir1Δ-1 lik1-5* lines #1 and #2 (Lanes 2-3). Lanes 4-6 are the respective negative controls where the reverse transcriptase was not included (-RT). *Elongation Factor 1α2* (*EF1A2*; At1g07930) was used as a positive control. *RKF1*= 472 bp, lanes 1,4:1ul cDNA used and 3ul PCR sample loaded, lanes 2,3,5,6:2ul cDNA used and 15ul PCR sample loaded (35 cycles for all lanes). *EF1A2*= 355bp, lanes 1-6:1ul cDNA used and 10ul PCR sample loaded (30 cycles for all lanes).

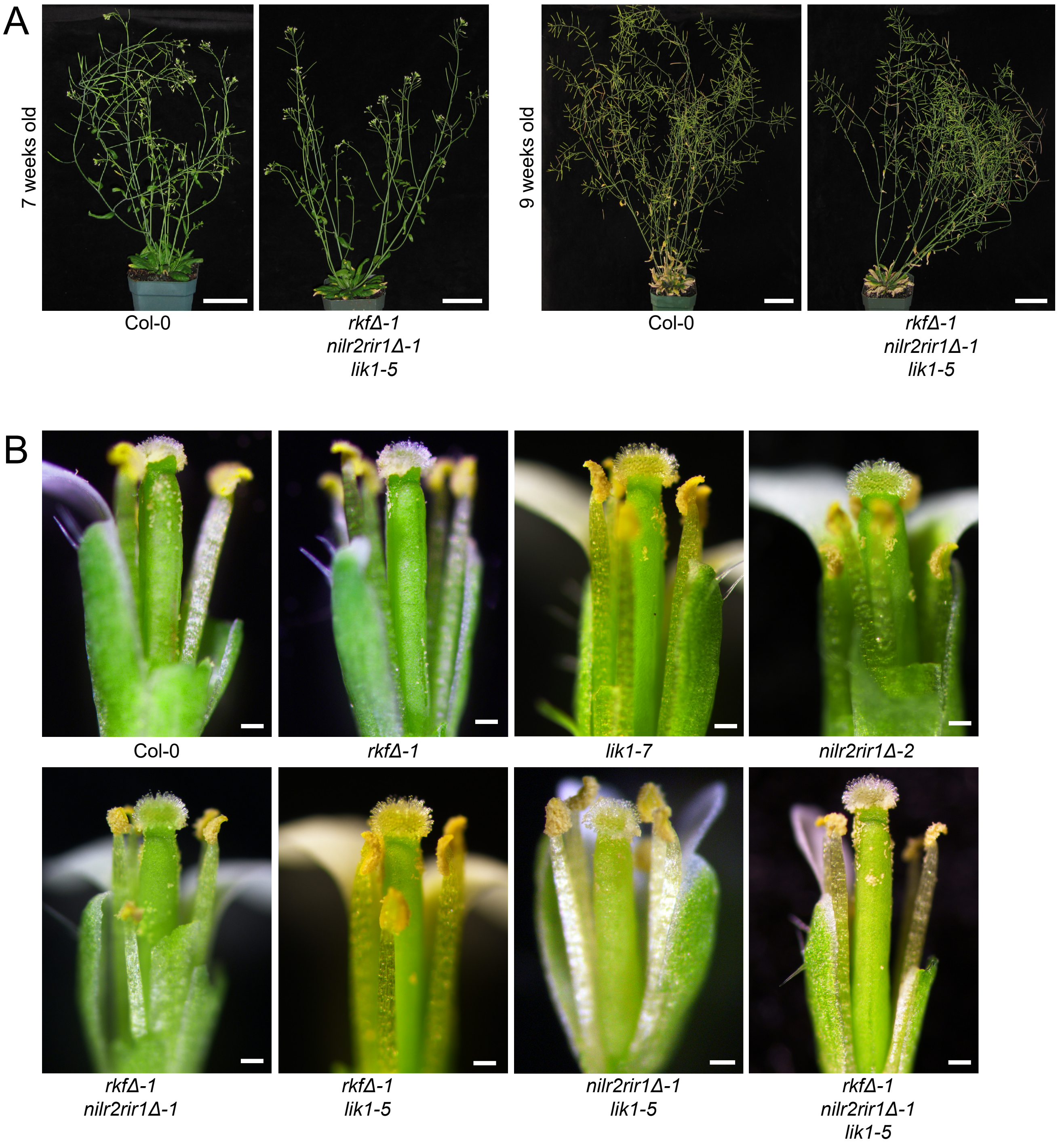


**Supplemental Figure 2. Representative images of Arabidopsis wild-type Col-0 and *LRR-MAL RK* mutant flowering plants and flowers.**

**A)** Whole plant photos were taken at 7 and 9 weeks. Scale bar = 5.0 cm.

**B)** Images of stage 14 flowers from *A. thaliana* wildtype Col-0 and the different *LRR-MAL RK* mutant combination plants (some petals and sepals have been removed to better show the pistil and anthers). Scale bar = 200 μm.

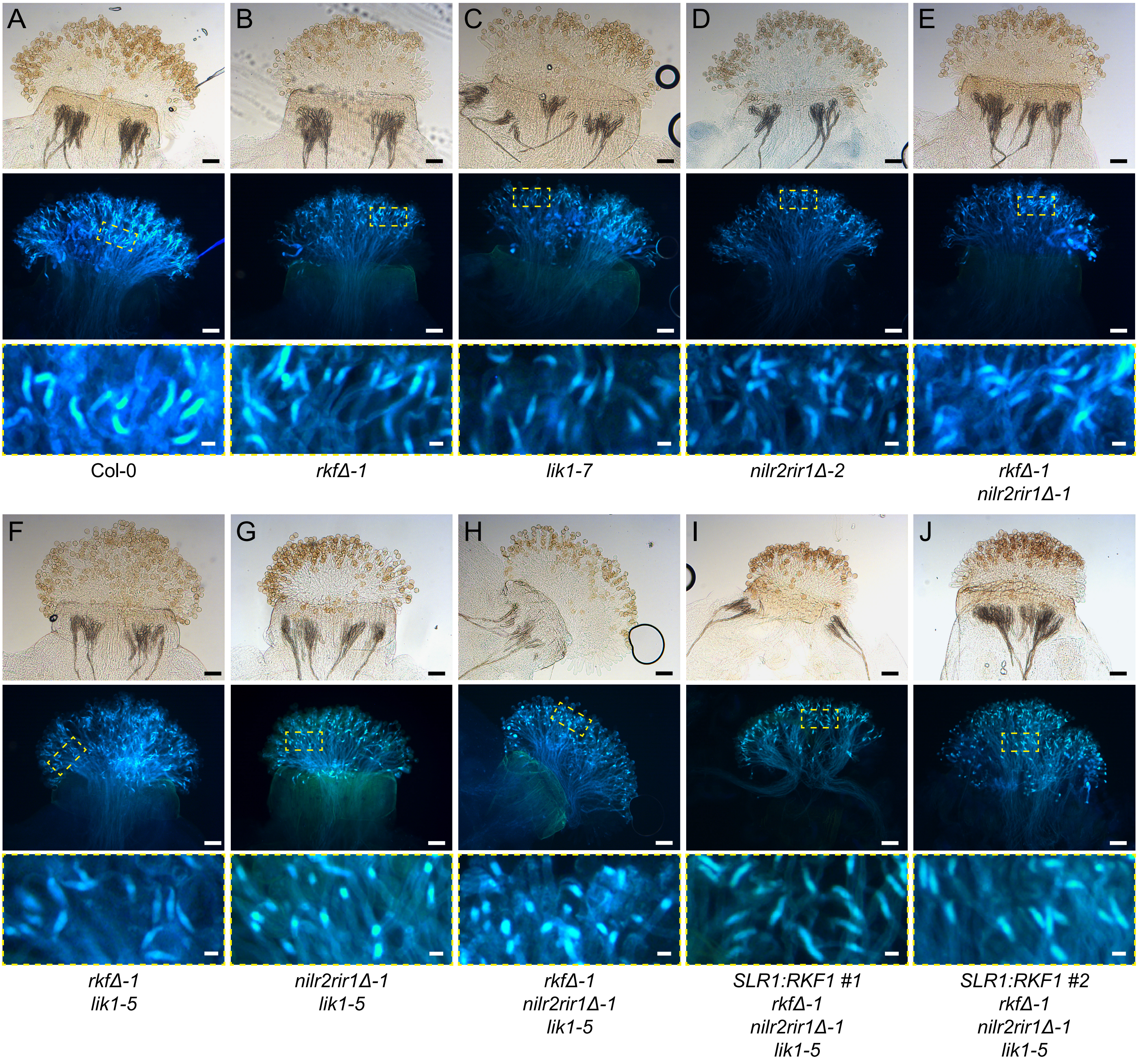


**Supplemental Figure 3. Representative images of aniline blue stained Col-0-pollinated pistils from *Arabidopsis* wildtype Col-0 and the *LRR-MAL RK* mutant plants.**
**A-J)** Stigmas from wildtype Col-0 and the different *LRR-MAL RK* mutant pistils were pollinated with Col-0 pollen and stained with aniline blue stained at 2-hours post-pollination. All plants were homozygous mutants for the indicated mutant alleles, and for each genotype, brightfield images are shown on the top and aniline blue stained images are shown on the bottom. For each set, the dashed yellow boxes in the middle panels outline the regions shown in bottom panels. Scale bar = 100 µm for the top two panels and 500 µm for the bottom panel. *SLR1:RKF1* is the stigma-expressed *RKF1* rescue construct.

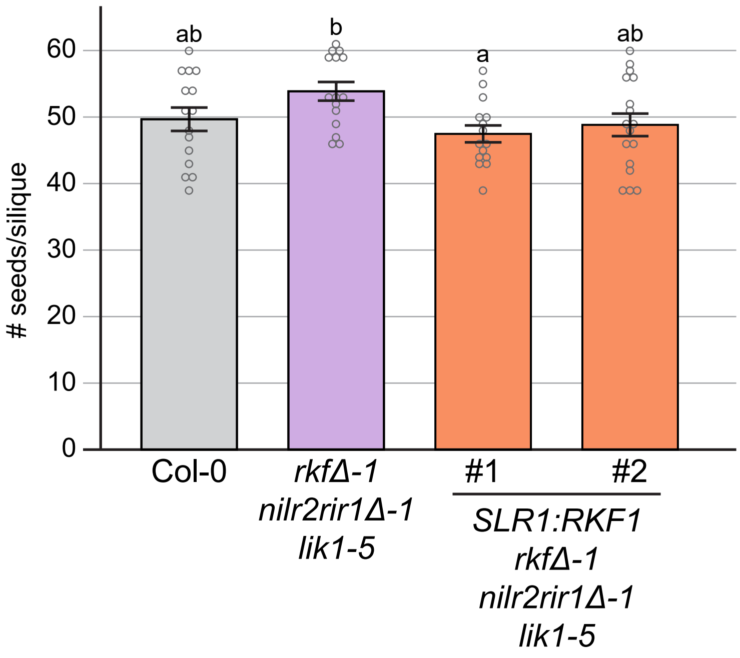


**Supplemental Figure 4. Seed set on naturally pollinated pistils.**
Data are shown as bar graphs of means ± SE with all data points displayed. n = 15-17 siliques per line. Letters represent statistically significant groupings of $P<0.05$ based on a one-way ANOVA with a Tukey-HSD post-hoc test.

**Supplemental Table S1. Expression profiles for *LRR-MAL RKs* in stigma transcriptome datasets**

| | | TRAVA RNA-Seq dataset *Klepikova et al 2016* http://travadb.org/ TMM-normalized raw values | | | |
|---|---|---|---|---|---|
| Name | AGI | Stigmas | Ovules | Carpels (ovules & stigmas removed) | Opened anthers, with pollen |
| BSR840 LIK1 | At3g14840 | 4451 | 2990 | 2280 | 466 |
| BSR440 RIR1 | At1g53440 | 1687 | 766 | 2364 | 82 |
| RKF1 | At1g29750 | 1056 | 631 | 1026 | 5289 |
| BSR650 | At1g07650 | 718 | 2372 | 2655 | 77 |
| RKFL1 | At1g29720 | 697 | 2 | 1 | 0 |
| BSR430 NILR2 | At1g53430 | 562 | 673 | 903 | 316 |
| CORK1 IGP1 | At1g56145 | 359 | 389 | 502 | 104 |
| IGP4 | At1g56140 | 249 | 407 | 417 | 153 |
| RKFL2 | At1g29730 | 62 | 62 | 60 | 110 |
| IGP2/3 | At1g56130 | 59 | 71 | 131 | 39 |
| | At1g56120 | 56 | 76 | 117 | 7 |
| RKFL3 | At1g29740 | 25 | 27 | 23 | 13 |
| | At1g53420 | 0 | 0 | 0 | 3 |

| Scale |
|---|
| 0 |
| 500 |
| 1000 |
| 1500 |
| 2000 |
| 2500 |
| 3000 |
| 3500 |
| 4000 |
| 4500 |
| >5000 |

| | | Stigmatic papillae RNA-Seq dataset *Gao et al 2018* | Pollen Coat-treated Stigma Microarray dataset *Iwano et al 2014* | | GUS activity *Wu et al 2016* | |
|---|---|---|---|---|---|---|
| Name | AGI | Stage 13 stigmatic papillae before pollination | 0 min Pollen Coat Average | 15 min Pollen Coat Average | Stigma | Style |
| BSR840 LIK1 | At3g14840 | 150 | 248 | 332 | Yes | Yes |
| BSR440 RIR1 | At1g53440 | 108 | 180 | 245 | Yes | Yes |
| RKF1 | At1g29750 | 39 | 128 | 144 | Yes | Yes |
| IGP4 | At1g56140 | 30 | 111 | 209 | No | No |
| CORK1 IGP1 | At1g56145 | 16 | 62 | 69 | No | No |
| BSR430 NILR2 | At1g53430 | 13 | 60 | 101 | Yes | Yes |
| BSR650 | At1g07650 | 7 | 124 | 149 | Yes | Yes |
| RKFL1 | At1g29720 | 6 | 38 | 37 | No | Weak |
| RKFL2 | At1g29730 | 4 | 42 | 39 | No | Weak |
| | At1g56120 | 2 | 25 | 26 | No | No |
| IGP2/3 | At1g56130 | 1 | 30 | 27 | Weak | Yes |
| RKFL3 | At1g29740 | 0 | 35 | 38 | No | No |
| | At1g53420 | 0 | 23 | 22 | Weak | Weak |

| Scale |
|---|
| 0 |
| 34 |
| 68 |
| 102 |
| 136 |
| 170 |
| 204 |
| 238 |
| 272 |
| 306 |
| >340 |

Heat maps generated with the BAR Heat mapper plus tool https://bar.utoronto.ca/

**Supplemental Table S2. Mutants used in this study.**

| Background | CRISPR deletion mutation | # Genes deleted | Genes | | | | Deletion Size | Reference |
|---|---|---|---|---|---|---|---|---|
| Col-0 | *rkfΔ-1* | 4 | *RKFL1* At1g29720 | *RKFL2* At1g29730 | *RKFL3* At1g29740 | *RKF1* At1g29750 | 24.5 kb | *Lee and Goring, 2021* |
| Col-0 | *lik1-7* | 1 | *LIK1* At3g14840 | | | | 5 kb | This paper: Supplementary Fig. 1 |
| Col-0 | *nilr2rir1Δ-2* | 2 | *NILR2* At1g53430 | *RIR1* At1g53440 | | | 14.9 kb | This paper: Supplementary Fig. 1 |
| *rkfΔ-1* | *lik1-5* | 1 | *LIK1* At3g14840 | | | | 5 kb | This paper: Supplementary Fig. 1 |
| *rkfΔ-1* | *nilr2rir1Δ-1* | 2 | *NILR2* At1g53430 | *RIR1* At1g53440 | | | 14.9 kb | This paper: Supplementary Fig. 1 |

| Higher Order Mutants | # Genes mutated | Genes | | | | | | |
|---|---|---|---|---|---|---|---|---|
| *nilr2rir1Δ-1* *lik1-5* | 3 | *NILR2* | *RIR1* | *LIK1* | | | | |
| *rkfΔ-1* *lik1-5* | 5 | *RKFL1* | *RKFL2* | *RKFL3* | *RKF1* | *LIK1* | | |
| *rkfΔ-1* *nilr2rir1Δ-1* | 6 | *RKFL1* | *RKFL2* | *RKFL3* | *RKF1* | *NILR2* | *RIR1* | |
| *rkfΔ-1* *nilr2rir1Δ-1* *lik1-5* | 7 | *RKFL1* | *RKFL2* | *RKFL3* | *RKF1* | *NILR2* | *RIR1* | *LIK1* |
